# Hepatic AMPK activation in response to dynamic REDOX balance is a biomarker of exercise to improve blood glucose control

**DOI:** 10.1101/2022.05.15.491995

**Authors:** Meiling Wu, Anda Zhao, Xingchen Yan, Hongyang Gao, Chunwang Zhang, Xiaomin Liu, Qiwen Luo, Feizhou Xie, Shanlin Liu, Dongyun Shi

**Affiliations:** Department of Biochemistry and Molecular Biology, School of Basic Medical Sciences, Fudan University, Shanghai, 200032, People’s Republic of China; Institute of Electronmicroscopy, School of Basic Medical Sciences, Fudan University, Shanghai, 200032, People’s Republic of China; Changning Maternity and Infant Health Hospital, East China Normal University, Shanghai, 200032, People’s Republic of China; Free Radical Regulation and Application Research Center of Fudan University, Shanghai, 200032, People’s Republic of China

**Author notes:** Correspondence to Prof. Dongyun Shi and Prof. Shanlin Liu, Department of Biochemistry and Molecular Biology, School of Basic Medical Sciences, Fudan University, NO.130 Dong’an Road, Shanghai, 200032, People’s Republic of China. Co-first authors contributed equally to this work (Meiling Wu, Anda Zhao, Xingchen Yan, Hongyang Gao).

**Keywords:** Redox balance, AMPK, exercise, glutathionylation

## Abstract

Antioxidant intervention is considered to inhibit reactive oxygen species (ROS) and alleviates hyperglycemia. Paradoxically, moderate exercise can produce ROS to improve diabetes. The exact redox mechanism of these two different approaches remains largely unclear. Here, by comparing exercise and antioxidants intervention on type 2 diabetic rats, we found moderate exercise upregulated compensatory antioxidant capability and reached a higher level of redox balance in the liver. In contrast, antioxidant intervention achieved a low-level redox balance by inhibiting oxidative stress. Both of these two interventions could promote glycolysis and aerobic oxidation mediated by hepatic AMPK activation, ameliorating diabetes. During exercise, different levels of ROS generated by exercise have differential regulations on the activity and expression of hepatic AMPK. Moderate exercise-derived ROS promoted hepatic AMPK glutathionylation activation. However, excess exercise increased oxidative damage, and inhibited the activity and expression of AMPK. Overall, our results illustrate that both exercise and antioxidant intervention improve blood glucose in diabetes by promoting redox balance, despite the levels of redox balance are different. Moreover, the activation and expression of AMPK could act as a biomarker to reflect the effective treatment range for diabetes. This finding provides theoretical evidence for the precise regulation of diabetes by antioxidants and exercise.

## Introduction

Diabetes mellitus is a chronic metabolic disorder disease, which has emerged as a global public health problem. According to the latest epidemiological data from the International Diabetes Federation, about 8.8% of the world population, or 415 million people, have diabetes as of 2019, about a 2.4% increase from that of 2010 (6.4%) (*1*). With the development of genomics, proteomics and metabolomics, it has been discovered by many studies that type 2 diabetes is associated with irreversible risk factors such as age, genetics, race, and ethnicity and reversible factors such as diet, physical activity and lifestyle (*2, 3*). Given the essential function of aerobic metabolism in glucose oxidation, mitochondrial damage and oxidative stress have been considered to play a critical role in the occurrence and development of diabetes (*4*). Exercise and antioxidant supplements are often suggested as essential therapeutic strategies in the early stages of type 2 diabetes (*5, 6*), with different mechanisms. It has been reported that chronic exercise training can alleviate oxidative stress and diabetic symptoms by improving cellular mitochondrial function and biogenesis in the diabetic state (*7*). Contradictorily, exercise also increases ROS production, while prolonged or high-intensity exercise could result in mitochondrial functional impairment to aggravate complications of diabetes (*8*). Since the 1970s, studies have demonstrated that 1 hour of moderate endurance exercise can increase lipid peroxidation in humans (*9, 10*). In 1998, Ashton directly detected increasing free radical levels in exercising humans using electron paramagnetic resonance spectroscopy (EPR) and spin capture (*11*). These results led to a great deal of interest in the role of ROS in physical exercise (*12-14*). Regarding the contradiction of exercise on ROS scavenging or production, James D Watson also hypothesized that type 2 diabetes is accelerated by insufficient oxidative stress rather than oxidative stress (*15*), based on the effect of exercise on diabetes treatment. Although Watson’s opinions supported that exercise could treat diabetes by producing ROS, whether exercise-induced ROS production is beneficial or detrimental to diabetes is still being debated. The specific regulation of ROS produced by exercise on diabetic blood glucose in *vivo* is unclear. In contrast, the general view of the antioxidant treatment for diabetes is that antioxidants reduce cytotoxic ROS and oxidation products, thus alleviating diabetes and achieving glycemic control (*16*). Our previous study also found that hepatic mitochondrial ROS scavengers and antioxidant substances inhibited the oxidative products such as MDA and 4-HNE in diabetic animals and favored glycemic control (*17, 18*). Exercise-induced oxidation and antioxidant administration, as two opposite approaches, could achieve the regulation of diabetes, respectively. However, the differences in redox mechanisms between these two approaches to diabetes treatment have not been fully understood.

It is well established that the increase of skeletal muscle glucose uptake during exercise is crucial in glycemic control (*19-21*). Considering that liver is another vital organ for maintaining blood glucose homeostasis, including storing, utilizing and producing glucose, exercise-induced hepatic redox metabolism is also significant. The activation of hepatic AMP-activated protein kinase (AMPK), which acts as a ‘metabolic master switch’, alleviates diabetes symptoms by reducing glycogen synthesis, increasing glycolysis, and promoting glucose absorption in surrounding tissues (*22*). Therefore, the activation of AMPK in the liver is significant for regulating glucose and lipid metabolism in the blood. Zmijewski et al. found that AMPK could be activated by hydrogen peroxide stimulation through direct oxidative modification (*23*). In contrast, other studies suggested that oxidative stress could disrupt the activation of the AMPK signaling pathway (*24, 25*). Our previous study explored the mechanism by which redox status contributes to hepatic AMPK dynamic activation. Under a low ROS microenvironment, GRXs mediated S-glutathione modification activates AMPK to improve glucose utilization. Meanwhile, under an excessive ROS microenvironment, sustained high level ROS might cause loss of AMPK protein (*26*). These studies indicate that oxidative modification can directly regulate AMPK activity in liver cells, thus activating downstream signaling pathways to regulate glucose and lipid metabolism. However, it is unclear why both antioxidant intervention and ROS produced by exercise can promote the seemingly contradictory phenomenon of AMPK activation. Moderate exercise has been proved significantly elevate systemic oxidative stress. At the same time, endogenous antioxidant defences also increased to counteract increased levels of ROS induced by exercise, leading to a higher level of redox balance(*27*). Thus, we hypothesized that both antioxidants and exercise could reach either high-level or low-level redox balance in diabetic individuals. Moreover, the activity and expression of AMPK might be a marker of redox balance *in vivo*.

Hence, the present study was designed to understand the different mechanisms of exercise and antioxidant intervention in diabetes and verify the activation of hepatic AMPK as a hallmark of dynamic redox balance. Firstly, we utilized the streptozotocin-high fat diet (STZ-HFD) induced type 2 diabetic (T2DM) model in rats to clarify the hepatic redox status in T2DM rats after the exercise or antioxidant intervention. Then, according to the exercise intensity and mode, we divided the exercise groups into three modes and found that AMPK activation could serve as a biomarker of redox balance and moderate exercise in diabetic treatment. Taken together, in this study, we found that AMPK activation and expression could reflect the threshold of exercise or antioxidant administration for diabetes treatment. It provides a theoretical basis for the precise regulation of diabetes by antioxidants and exercise.

## Results

### 1. Execise promotes antioxidant levels through producing ROS, leading to a high level of REDOX balance in the liver

To investigate the hepatic redox regulation in diabetes after exercise intervention, we established the T2DM rat model by feeding HFD followed by a low dose of STZ injection (35 mg/kg). The exercise intervention was started from Day 0 to Day 28 (Fig. 1A). According to previous studies, the initial speed of exercise was 15 m/min, and the speed was increased by 3 m/min every 5 min. After the speed reached 20 m/min, the speed was maintained for another 60 min with slope of 5%. The exercise intensity was 64%-76% VO_2max_ (*28*). The low-intensity continuous exercise (CE) can be regarded as aerobic exercise.

**Fig. 1.**
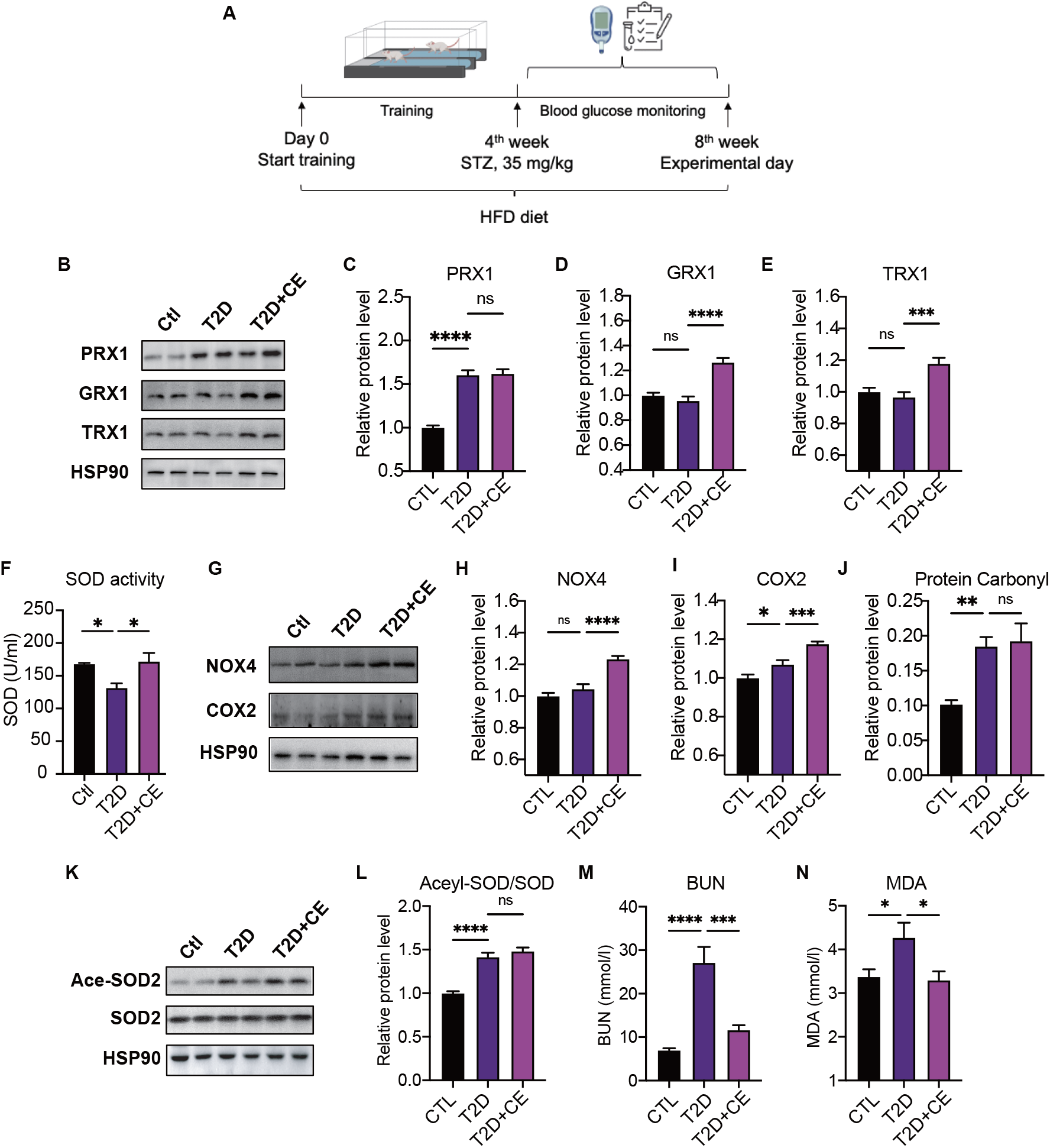
Exercise induced ROS production in exercise group and increased the antioxidant status. **A**. Experimental design. T2DM rats model was fed by high-fat diet plus a low dose of STZ injection (35 mg/kg). The high-fat diet (HFD, 60% calories from fat) was started from the 1^st^ week to the 8^th^ week. The exercise intervention was started from 1^st^ week to 4^th^ week. **B-E**. Representative protein level and quantitative analysis of PRX1 (27 kDa), Grx1 (17 kDa), Trx1 (12 kDa) and HSP90 (90 kDa) in the rats in the Ctl, T2D and T2D + CE groups. The rat livers were homogenized by 1% SDS and analyzed by Western blots with the appropriate antibodies. **F-I**. Representative protein level and quantitative analysis of NOX4 (27 kDa), COX2 (17 kDa) and HSP90 (90 kDa) in the rats in the Ctl, T2D and T2D + CE groups. **J-K**. Representative protein level and quantitative analysis of Ace-SOD2 (27 kDa), SOD2 (17 kDa) and HSP90 (90 kDa) in the rat in the Ctl, T2D and T2D + CE groups. **L-M**. Liver MDA content (L) and BUN (M) level was detected in the rats of Ctl, T2D and T2D + CE groups. (ns: not significant; *P < 0.05, **P < 0.01, ***P < 0.001, ****P < 0.0001 compared with all groups by one-way ANOVA and Tukey’s post hoc test; data are expressed as the mean ± SEM; n = 4-8 per group).

Firstly, we detect the expression of antioxidant enzymes and oxidase in liver tissue. As redox proteins regulate the redox state *in vivo*, the protein expression of GRX and TRX were found to be up-regulated during exercise intervention (Fig. 1B-E). Notably, the PRX expression also showed a trend of increase (Fig. 1B-C). Parallelly, the expressions of NADPH oxidase 4 (NOX4) and cyclooxygenase 2 (COX2) in the liver were also significantly up-regulated in the exercise group (Fig. 1F-H). Protein carbonylation is a type of protein oxidation that can be promoted by ROS. However, we found that the exercise group did not decrease the protein carbonylation level (Fig. 1I). As shown in Fig. 1J-K, the acetylation level of MnSOD also shows an increase in the exercise group, indicating the inactivation of mitochondrial MnSOD. MDA, a biomarker of lipid peroxidation, was also significantly up-regulated in the diabetic group but decreased in exercise group (Fig.1L). Meanwhile, specific markers related to kidney dysfunction, such as the blood urea nitrogen (BUN) level, were also significantly increased in the diabetic rat group. Exercise intervention reduced the BUN level (Fig.1M). These results indicated that the high ROS production in the exercise group could compensatory increase the antioxidant status to avoid oxidative damage. It suggests that exercise can promote redox to reach a high level of balance, therefore ROS produced by exercise does not lead to oxidative damage.

### 2. Antioxidant intervention alleviates blood glucose through reducing oxidative stress, leading to a low level of REDOX balance in the liver

Recent studies have suggested that NADPH oxidase is one of the primary sources of ROS (*29*). Apocynin has already been characterized as an NADPH oxidase inhibitor in the early 1980s, and it can also act as an antioxidant (*30*). Our previous study showed that apocynin intervention alleviated blood glucose by inhibiting oxidative products. Compared with the exercise intervention, the antioxidant intervention was also started from Day 0 to D28 in this study (Fig. 2A). We found that apocynin intervention decreased the protein carbonylation level and MDA level in the liver (Fig. 2B-C). Also, the TAOC level increased after apocynin treatment (Fig. 2D). The random blood glucose and oral glucose tolerance (2 h after oral glucose, OGTT) decreased in the apocynin intervention group compared with the diabetic rat group (Fig. 2F-G). Consistent with the apocynin intervention group, the exercise group also showed lower random blood glucose levels and 2h OGTT (Fig. 2H-I). These studies indicated that the apocynin treatment inhibited the protein oxidative damage and alleviated blood glucose.

**Fig. 2.**
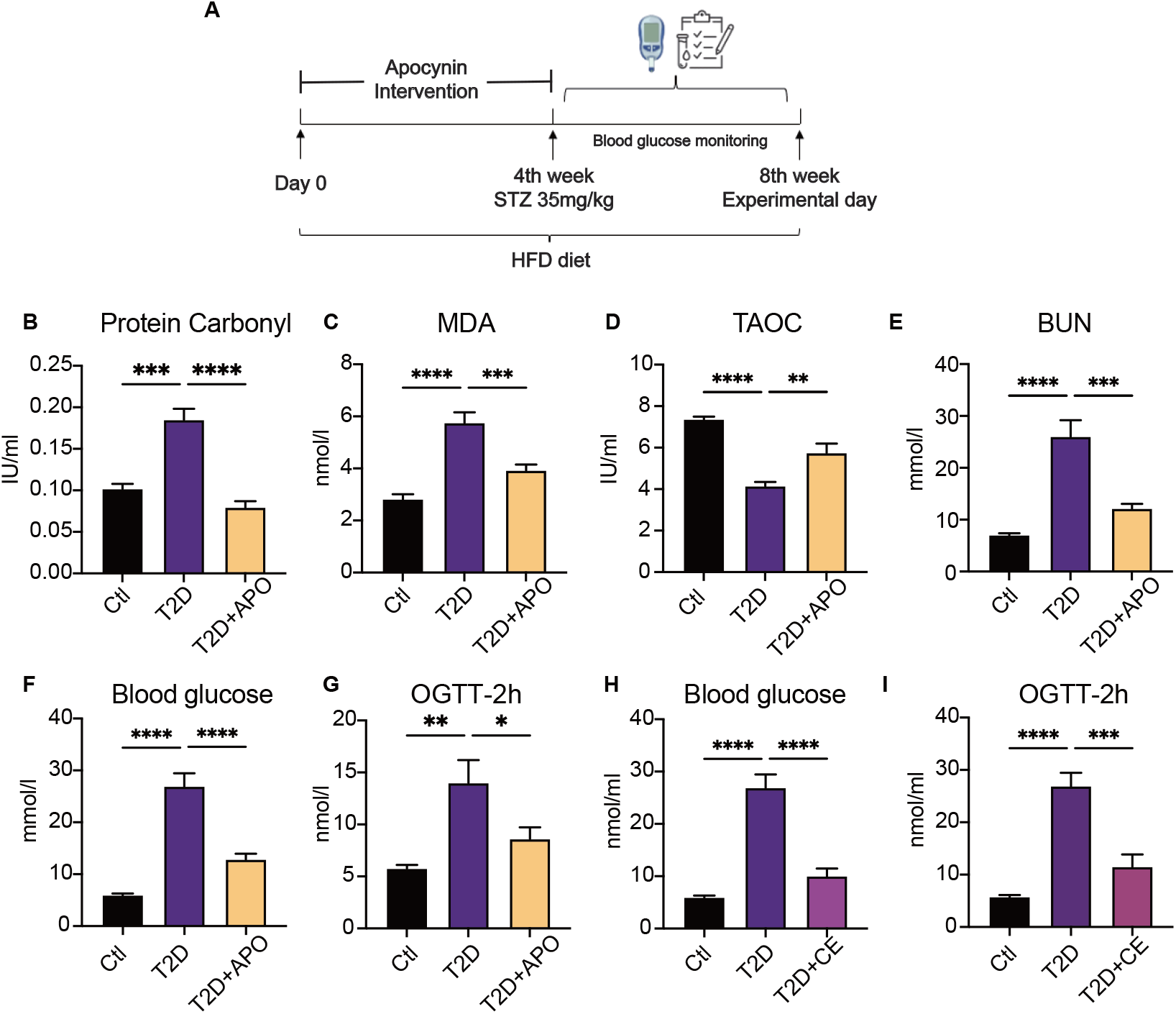
Antioxidant intervention alleviates blood glucose through promoting the upregulation of reducing levels. **A**. Experimental design. T2DM rats model was fed by high-fat diet plus a low dose of STZ injection (35 mg/kg). The apocynin intervention was started from 1^st^ week to 4^th^ week. **B**. Liver protein carbonylation was detected in the rats of Ctl, T2D and T2D + Apocynin (APO) groups. **C-E**. Liver MDA content (C), TAOC (D) and BUN (E) level was detected in the rats of Ctl, T2D and T2D + APO groups. **F**. Postprandial blood glucose levels of Ctl, T2D and T2D + APO groups at the end of 8^th^ week. **G**. Blood glucose level after oral glucose administration in Ctl, T2D and T2D + APO groups at the end of 8^th^ week. **H**. Postprandial blood glucose levels of Ctl, T2D and T2D + CE groups at the end of 8^th^ week. **I**. Blood glucose level after oral glucose administration in Ctl, T2D and T2D + CE groups at the end of 8^th^ week (*P < 0.05, **P < 0.01, ***P < 0.001, ****P < 0.0001 compared with all groups by one-way ANOVA and Tukey’s post hoc test; data are expressed as the mean ± SEM; n

### 3. Moderate exercise-generated ROS production promotes phosphorylated activation of AMPK and reduces blood glucose level, while excessive exercise-generated oxidative stress reduces AMPK expression and exacerbates diabetes

In order to find out the biomarkers that could reflect moderate exercise to improve blood glucose control, diabetic rats were divided into short-term continuous exercise (CE), intermittent exercise (IE), and excessive exercise (EE) according to the exercise intensity and mode (*28*). We found that the random blood glucose and 2h OGTT in CE and IE treated diabetic rats decreased (Fig. 2A-B). In contrast, EE intervention did not improve blood glucose but increased random and 2h OGTT, without statistical significance (Fig. 3A-B). Next, we detected the expression of antioxidant enzyme and oxidase in the liver tissue of exercise-treated T2D rats. Hepatic MDA concentration showed significant up-regulation in the diabetic group but a decrease in continuous and intermittent exercise (Fig.3D). We found that the CE group did not obviously change the protein carbonylation level. However, the EE intervention promoted the protein carbonylation in the liver, indicating the mode of action is not free radical scavenging but ROS production (Fig.3E).

**Fig. 3.**
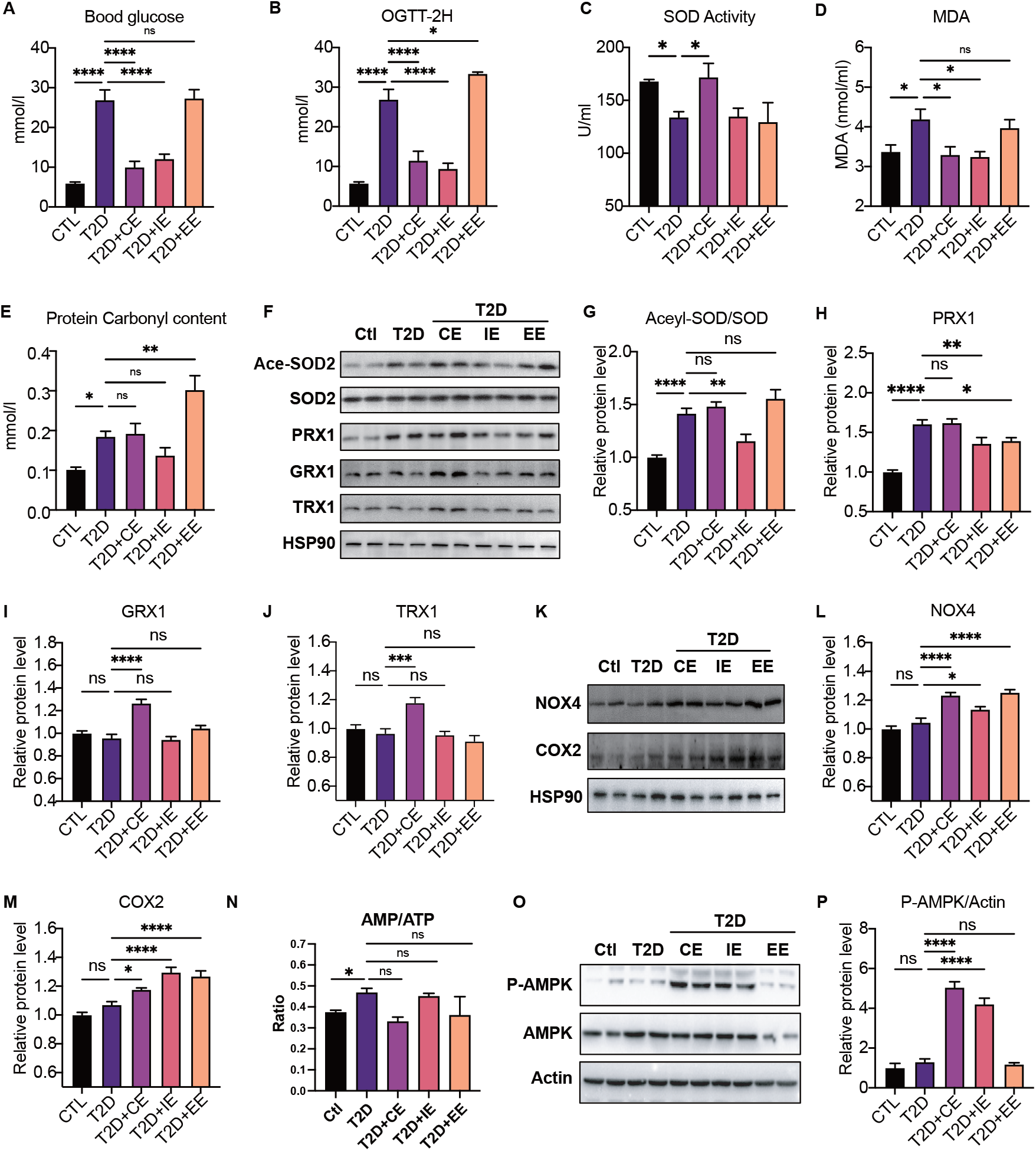
Moderate exercise-generated ROS production promotes phosphorylated activation of AMPK and reduces blood glucose level, while excessive exercise-generated oxidative stress reduces AMPK expression and exacerbates diabetes. **A**. Postprandial blood glucose levels of Ctl, T2D, T2D + CE, T2D + IE and T2D + EE groups at the end of 8^th^ week. **B**. Blood glucose level after oral glucose administration in Ctl, T2D, T2D + CE, T2D + IE and T2D + EE groups at the end of 8^th^ week. **C-E**. SOD activity (C), liver MDA content (D) and liver protein carbonylation content (E) level was detected in the rats of Ctl, T2D, T2D + CE, T2D + IE and T2D + EE groups. **F-J**. Representative protein level and quantitative analysis of Ace-SOD2 (27 kDa), SOD2 (17 kDa), PRX1 (27 kDa), Grx1 (17 kDa), Trx1 (12 kDa) and HSP90 (90 kDa) in the rats in the Ctl, T2D, T2D + CE, T2D + IE and T2D + EE groups. **K-N**. Representative protein level and quantitative analysis of NOX4 (27 kDa), COX2 (17 kDa) and HSP90 (90 kDa) in the rats in the Ctl, T2D, T2D + CE, T2D + IE and T2D + EE groups. **O-P**. Representative protein level and quantitative analysis of P-AMPK (67 kDa), AMPK (67 kDa) and Actin (45 kDa) in the rats in the Ctl, T2D, T2D + CE, T2D + IE and T2D + EE groups. (ns: not significant; *P < 0.05, **P < 0.01, ***P < 0.001, ****P < 0.0001 compared with all groups by one-way ANOVA and Tukey’s post hoc test; data are expressed as the mean ± SEM; n = 4-8 per group).

The ROS-generating NADPH oxidases (NOXs) have been recognized as one of the main sources of ROS production in cells (*31*). Cyclooxygenase 2 (COX2) activity could also act as a stimulus for ROS production (*32*). The increase of NOX4 and COX2 in the EE group indicated the highest oxidation level (Fig. 3K-L). As shown in Fig. 3F-G, although the acetylation level of MnSOD was found to increase significantly in the CE and EE group, which presented the inactivation of MnSOD, the antioxidant enzyme GRX and TRX were found to be up-regulated during CE intervention (Fig. 3I-J). Thus, the CE intervention maintains a high-level balance of redox state. Considering the decrease of antioxidant enzymes in the EE group, the REDOX balance in EE group was disrupted. Therefore, MDA in the CE group did not increase, while the increased MDA in the EE group indicated oxidative damage (Fig. 3D). Among these three exercise modes, the IE group showed the lowest level of oxidation (the minor increase in COX and a slight decrease in carbonylation). Although the levels of antioxidant enzymes such as GRX, TRX, and PRX did not increase, the activity of MnSOD also increased significantly (the level of acetylation decreased) (Fig. 3F). The reduction of MDA level also indicates IE group did not form oxidative damage (Fig. 3D), indicating the IE group could also maintain a relatively high level of redox balance.

Notably, the phosphorylation of AMPK showed different patterns in three kinds of exercise, among which both CE and IE intervention could promote the phosphorylation of AMPK compared to the diabetic rats (Fig. 3O-P). EE intervention did not increase the content of AMPK phosphorylation, which might be caused by the reduction of AMPK level. Meanwhile, the ratio of AMP to ATP was detected, and exercise-activated AMPK did not exhibit AMP-dependent characteristics at this time (Fig. 3N). These results suggested that moderate exercise-generated ROS may directly promote AMPK phosphorylation activation (independent of AMP upregulation) and reduce blood and liver glucose levels. However, excessive exercise-generated oxidative stress reduces AMPK expression and exacerbates diabetes.

### 4. Moderate exercise promoted glycolysis and mitochondrial tricarboxylic acid cycle in the liver of diabetic rats

Next, we further explored the mechanism by which inhibiting blood glucose during CE and IE intervention. Fructose-2,6-diphosphate (F-2,6-P2; also known as F-2,6-BP), which is a product of the bifunctional enzyme 6-phosphofructose 2-kinase/fructose 2,6-diphosphatase 2 (PFK/FBPase 2, also known as PFKFB2), is a potent regulator of glycolytic and gluconeogenic flux. The phospho-PFKFB2 to PFKFB2 ratio represents the glycolytic rate. A high ratio of phospho-PFKFB2:PFKFB2 leads to an increase in the F-2,6-P2 level and the allosteric activation of phosphor-fructose kinase 1 (PFK1), while a low ratio leads to a decrease in F-2,6-P2 and an increase in gluconeogenesis (*33*). The overexpression of bifunctional enzymes in mouse liver can reduce blood glucose levels by inhibiting hepatic glucose production (*34*). Therefore, bifunctional enzymes are also a potential target for reducing hepatic glucose production. In our study, the p-PFK2:PFK2 ratio decreased in the diabetic rats but was enhanced by CE and IE intervention (Fig. 4A-C), suggesting that CE and IE could reverse gluconeogenesis to glycolysis by enhancing PFK/FBPase. Meanwhile, the substrates of the glycolytic pathway (such as DHAP, Fig. 4D) and the tricarboxylic acid cycle (such as citrate, succinate and malate, Fig. 4D) showed an upward trend. These results illustrated that moderate exercise promoted glucose catabolism in the liver of diabetic rats (Fig. 4E).

**Fig. 4.**
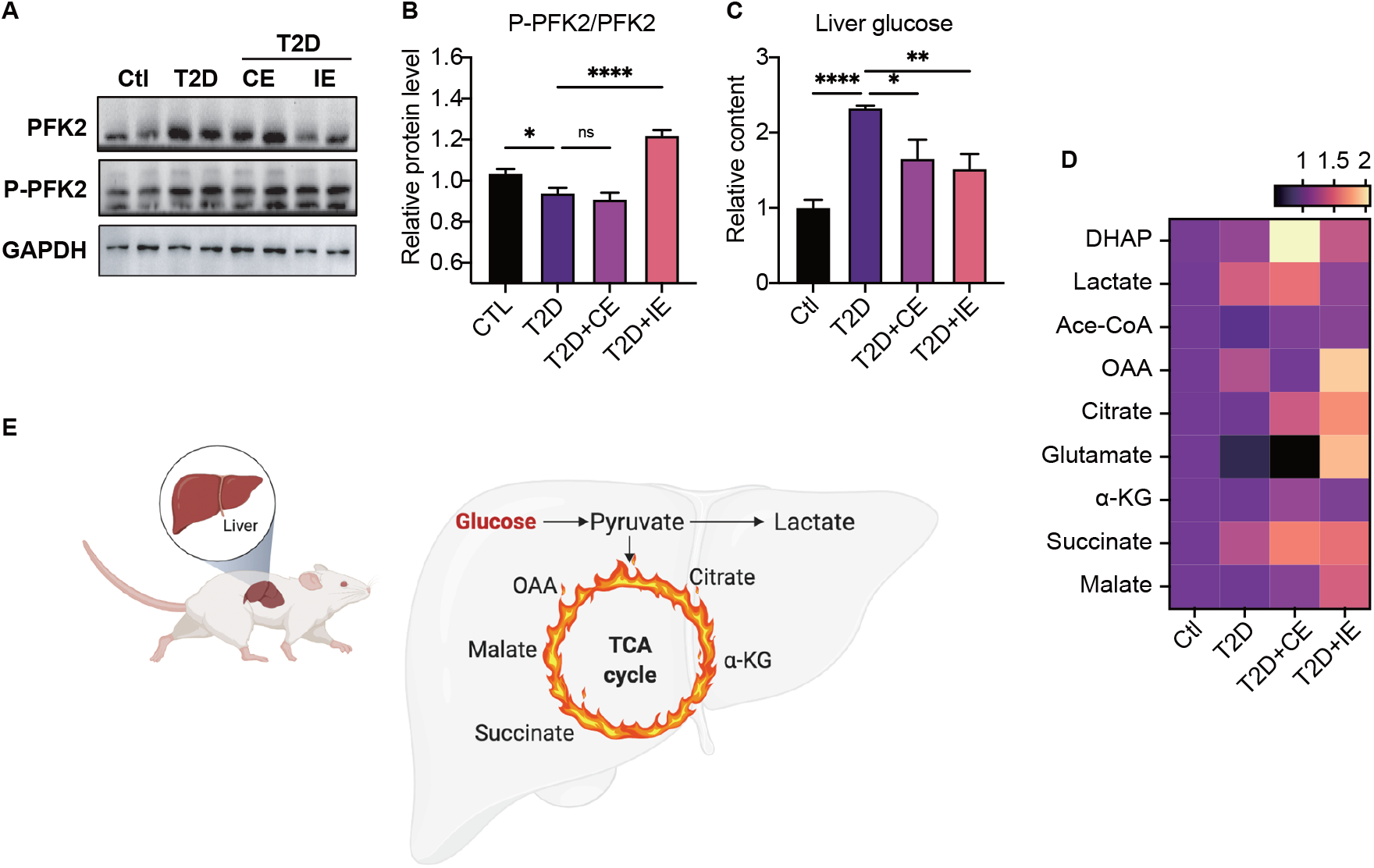
Moderate exercise promoted glycolysis and mitochondrial tricarboxylic acid cycle in the liver of diabetic rats. **A-B**. Representative protein level and quantitative analysis of P-PFK2 (64 kDa), PFK2 (64 kDa) and GAPDH (37 kDa) in the rats in the Ctl, T2D, T2D + CE and T2D + IE groups. **C**. Liver glucose level after oral glucose administration in Ctl, T2D, T2D + CE and T2D + IE groups at the end of 8^th^ week. **D**. Relative concentrations of substrates for glycolysis (DHAP and Lactate) and the tricarboxylic acid cycle (citrate, succinate and malate) in the rat of Ctl, T2D, T2D + CE and T2D + IE groups. The concentration of substrates was analyzed by LC-MS/MS. **E**. Schematic diagram illustrating the effect of CE and IE on glycolysis and mitochondrial tricarboxylic acid cycle (ns: not significant; *P < 0.05, **P < 0.01, ****P < 0.0001 compared with all groups by one-way ANOVA and Tukey’s post hoc test; data are expressed as the mean ± SEM; n = 6-8 per group).

### 5. Moderate exercise inhibited hepatic mitophagy, while excessive exercise promoted hepatic mitophagy and inhibited the mitochondrial biogenesis

The electron transport associated with the mitochondrial function is considered the major process leading to ROS production during exercise (*35*). To further explore the downstream signal of AMPK activation in moderate and excessive exercise, we detected the protein expression of mitochondrial dynamic and mitochondrial biogenesis. According to the result in Figure 5A-E, we found that the mitochondrial fusion protein MFN significantly decreased in the liver of the excessive exercise group, and the mitochondrial fission protein (Fis) and autophagy-related protein ATG5 and LC3B did not change, compared with the diabetic group. Notably, the ATG5 and LC3B levels decreased in the CE and IE group, compared with the diabetic group (Fig. 5A-E). PGC1α is a transcriptional coactivator, a central inducer of mitochondrial biogenesis in cells (*36*). The expression of PGC1α increased in the CE and IE group, but not in the EE group (Fig. 5F). These results indicated that moderate exercise ameliorated mitochondrial biogenesis and autophagy in the liver. However, excessive exercise aggravated mitochondrial fission and did not exhibit autophagy alleviation. Parallelly, the mitochondria structure of the live tissue in the EE group is fragmented and showed greatly diminished cristae and swelling matrix under transmission electron microscopy, which is functionally reflected as a defect in oxidative phosphorylation. However, the CE and IE group showed increased numbers of cristae and a clear structure of mitochondrial cristae (Fig. 5H). These results all show that the *in vivo* mitochondrial ROS burst caused by excessive exercise inhibits the expression of AMPK and promotes mitophagy. The damage to the mitochondrial dynamics and structure in liver tissue leads to abnormal aerobic oxidation, thereby aggravating diabetes

**Fig. 5.**
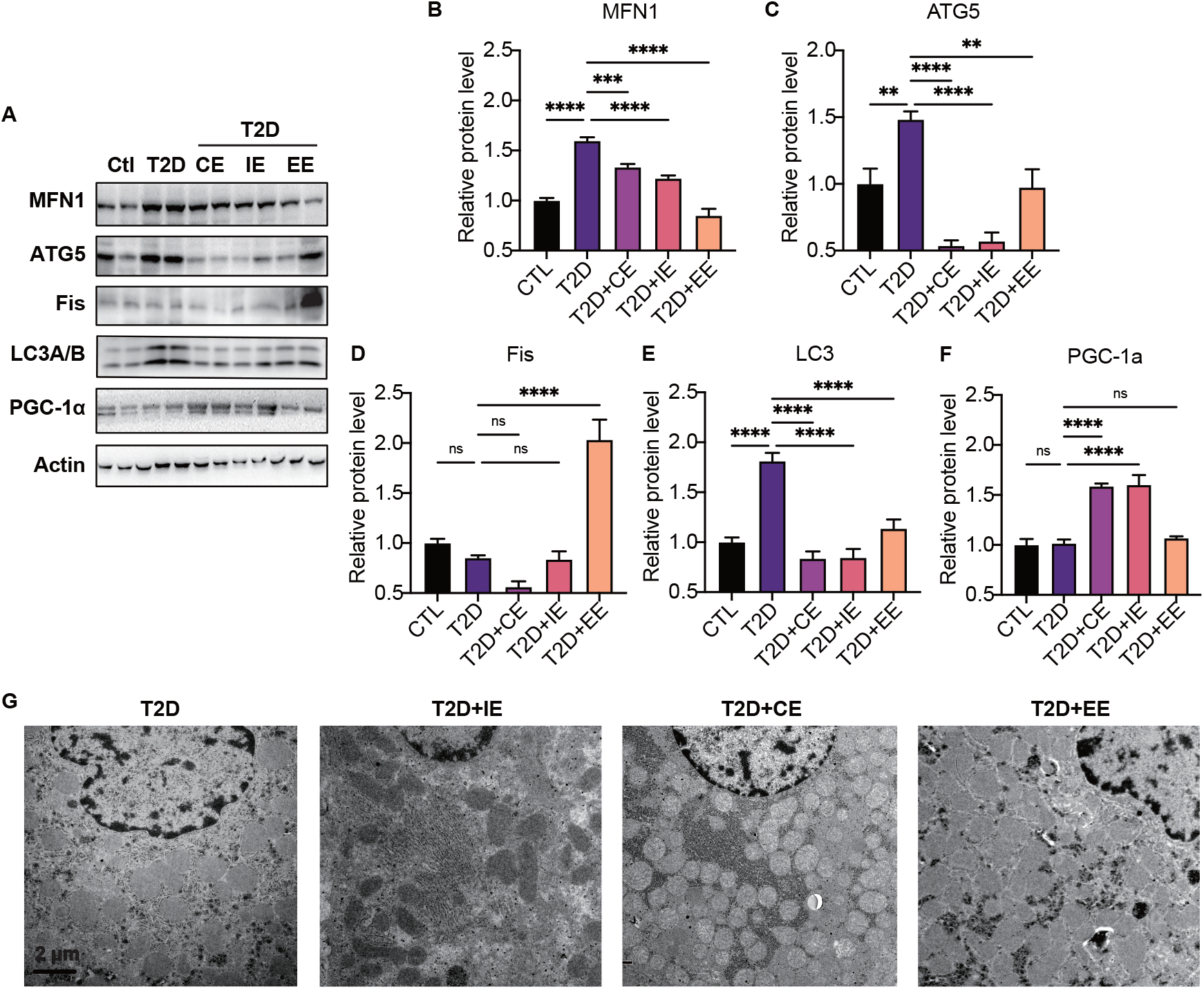
Excessive exercise promoted hepatic mitophagy, inhibited the mitochondrial biogenesis and destroyed the mitochondrial structure. **A-F**. Representative protein level and quantitative analysis of MFN1 (82 kDa), ATG5 (55 kDa), FIS (25 kDa), LC3A/B (14,16 kDa), PGC-1α (130 kDa) and Actin (45 kDa) in the rats in the Ctl, T2D, T2D + CE, T2D + IE and T2D + EE groups. **G**. TEM analysis of the ultrastructure of hepatocytes in the rats in the T2D, T2D + CE, T2D + IE and T2D + EE groups. (Scale bar = 2 μm; ns: not significant; *P < 0.05, **P < 0.01, ***P < 0.001, ****P < 0.0001 compared with all groups by one-way ANOVA and Tukey’s post hoc test; data are expressed as the mean ± SEM; n = 8 per group).

### 6. ROS differentially regulated AMPK activation through GRX-mediated glutathionylation within redox balance threshold

To further elucidate the relationship between redox balance and AMPK activation in cellular environment, L02 cells were intervened with different concentrations of H_2_O_2_ (50-200 μmol/l) to mimic the different ROS levels *in vivo* (Fig. 6A). We found the expression of 3-NT and 4-HNE increased after H_2_O_2_ intervention (100-200 μmol/l, Fig. 6B). The acetylation level of Mn-SOD also increased after H_2_O_2_ intervention (Fig. 6C). Significantly, oxidative stress marker 3-nitrotyrosine (3-NT) level highly increased at the concentration of 200 μmol/l, indicating the H_2_O_2_ intervention at 200 μmol/l exceeded the threshold of redox balance, thus causing oxidative damage. Our previous study shows that optimal ROS would activate AMPK through GRX-mediated S-glutathionylation (*26*). As shown in Fig. 6D-F, exposure to 50 and 100 μmol/l H_2_O_2_ led to an increase of GSS-protein adduct, concomitant with the AMPK phosphorylation and glutathionylation (Fig. 6E-F), suggesting that the ROS level within redox balance threshold could induce glutathionylation and phosphorylation of AMPK. However, when the concentration of ROS was too high to exceed the redox balance threshold, the AMPK protein would be partially degraded, thereby inhibiting its activity. Exposure to 200 μmol/l H_2_O_2_ led to a decrease in AMPK glutathionylation, phosphorylation as well as protein content (Fig. 6D-F). These results indicated that the increased ROS within the redox balance threshold could promote the AMPK activation through GRX-mediated S-glutathionylation.

**Fig. 6.**
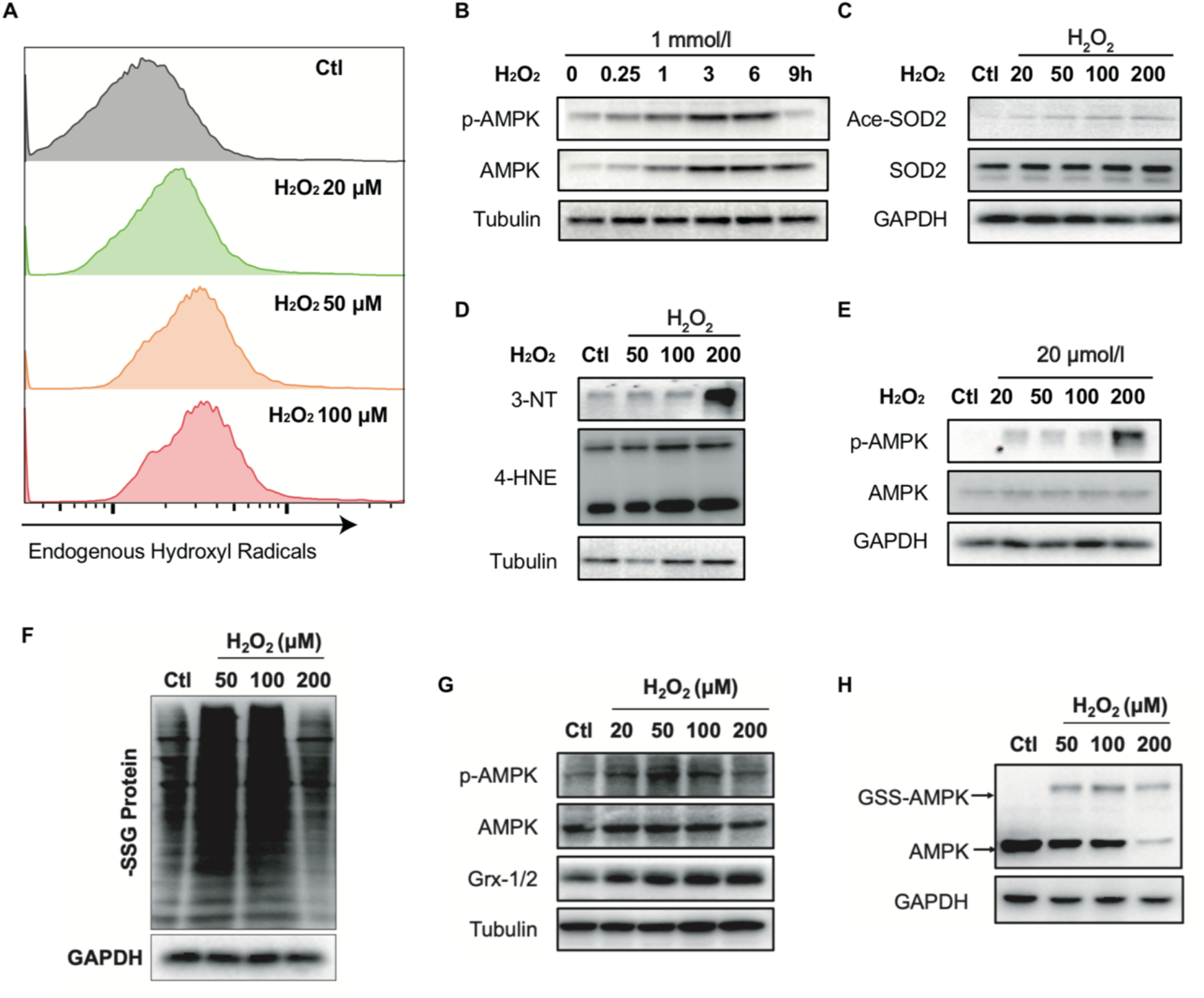
ROS differentially regulated AMPK activation through Grx-mediated glutathionylation within redox balance threshold. **A**. Analysis of superoxide generation using flow cytometry and a hydroethidine probe in L02 cells under H2O2 stress (50–200 μmol/l, 30 min). **B**. Representative protein level of 4-HNE, 3-NT and GAPDH (37 kDa) in L02 cells under H2O2 stress (50–200 μmol/l, 30 min). **C**. Representative protein level of Aceyl-SOD2, SOD2 and GAPDH (37 kDa) in L02 cells under H2O2 stress (50–200 μmol/l, 30 min). **D**. GSS-adduct protein level in L02 cells under H2O2 stress (50–200 μmol/l). L02 cells loaded with EE-GSH-biotin were incubated with/without H2O2 for 30 min, and the amounts of GSS-protein adduct formation were determined using non-reducing SDS-PAGE and Western blot analysis with streptavidin-HRP. **E**. Representative protein level of P-AMPK (67 kDa), AMPK (67 kDa), Grx-1/2 (12 kDa) and Tubulin (55 kDa) in L02 cells under H2O2 stress (50–200 μmol/l, 30 min). **F**. AMPK cysteine gel shift immunoblot. Cysteine dependent shifts by incubation of AMPK protein with glutathione reductase and PEG-Mal. PEG2-mal labelled glutathionylation modification shifts AMPK by ∼10 kDa above the native molecular weight.

At the same time, we also detected the substrates of glycolysis and aerobic oxidation at different concentrations of H_2_O_2_. We found the exposure to 20-100 μmol/l H_2_O_2_, which made cells within the redox balance threshold, showed a trend of increase on glycolysis and aerobic oxidation substrates, indicating the increase of hepatic glucose catabolism (data not shown).

## Discussion

It is known that both antioxidants and exercise can substantially benefit and ameliorate hyperglycaemia through ROS-mediated mechanisms in diabetes patients. However, antioxidant intervention reduces oxidative stress, while exercise produces ROS. The differences in redox mechanism between these two opposite approaches to diabetes treatment have not been fully explored. Therefore, it is still unclear how to use exercise and antioxidants scientifically and rationally to treat diabetes effectively.

The remission of diabetes by antioxidant intervention has been well documented. Some compounds in food that have a vigorous antioxidant activity or inhibit NADPH oxidase, such as polyphenols and flavonoids (*37*), have been confirmed to improve blood glucose and relieve type 2 diabetes in animal experiments. Several clinical trials also demonstrated the relief of antioxidants in diabetic hyperglycaemia (*38, 39*). Our previous study found hepatic mitochondrial ROS scavenger and antioxidant substances inhibited the oxidative products such as MDA and 4-HNE in diabetic mice and rats and benefitted blood glucose control (*18*). These evidence indicates that reducing the oxidative level of diabetic animals could treat diabetes. Hepatic AMPK regulates cellular and whole-body energy homeostasis, signals to stimulate glucose uptake in skeletal muscles, fatty acid oxidation in adipose (and other) tissues, and reduces hepatic glucose production (*40, 41*). Numerous pharmacological agents (including the first-line oral drug metformin), natural compounds, and hormones are known to activate AMPK (*42-45*). Moreover, our previous found that antioxidant (apocynin) intervention in diabetic rats could promote the phosphorylation and activation of AMPK protein, thereby regulating hepatic glucose metabolism (*46*). Taken together, we found that the activation of AMPK by antioxidant intervention was accompanied by a decrease in oxidative stress level in diabetic rats, resulting in a low level of redox balance to benefit diabetic hyperglycaemia.

Many medical research and public health recommendations support regular exercise to mitigate symptoms of many diseases, including psychiatric, neurological, metabolic, cardiovascular, pulmonary, musculoskeletal, and even cancer (*47*). John Holloszy’s studies found that exercise improved insulin sensitivity in patients with type 2 diabetes and provided a better understanding of how muscle adapts to endurance exercise (*19-21, 48-50*). Although the benefits of exercise are irrefutable, excessive exercise is harmful (*8*). It is unclear how to control the amount and intensity of exercise. Recently, Chrysovalantou et al. found that NADPH oxidase 4 (NOX4) is a crucial protein of exercise to regulate adaptive responses and prevent insulin resistance (*51*). We found that exercise can indeed increase NOX expression, but NOX is also upregulated in excessive exercise. Although Chrysovalantou’s research puts more emphasis on the role of the redox environment in exercise, the biomarkers of moderate exercise for diabetes remain uncertain. In this study, according to the exercise intensity and mode, we divided the exercise groups into three modes: CE, IE and EE. We found that moderate exercise (including CE and IE) promoted hepatic ROS production and up-regulated a compensatory increase in antioxidant capability, forming a higher level balance of redox state. However, excessive exercise increases mitochondrial oxidative stress level and cause oxidative damage, which is not beneficial for glycaemic control in diabetes.

During exercise, the activation of AMPK in skeletal muscle was considered mainly caused by the increase of intracellular AMP:ATP ratio and phosphorylation of Thr172 on the “activation loop”7 of the α-subunit (*52*). The activation of AMPK leads to the inhibition of mTORC1 activity and activation of PGC-1α, thereby enhancing mitochondrial biogenesis (*53*). Early studies clarified that the activation of AMPK is related to the liver energy state during exercise (*54*). There are some papers against the role of AMPK in regulating glucose uptake during exercise (*55-57*), but these studies mainly focused on skeletal muscle glucose uptake but not the liver. Although the glucose was uptake by primarily skeletal muscle after exercise, the liver is prominent in whole-body glucose tolerance (*58*). Increasing evidence showed that AMPK is a redox-sensitive protein, and its cysteine 299 and 304 sites are likely to be regulated by the oxidation of hydrogen peroxide (*23, 59, 60*). Thus, the phosphorylation of AMPK might be directly activated through ROS regulation during exercise, not only depending on the increase of AMP. Our previous study found that both oxidation and reduction can promote AMPK activation (*26*), actually, this is because the redox balance state induced by oxidation or reduction intervention activates AMPK. In this study, we demonstrates that both antioxidant intervention and moderate exercise can activate AMPK oxidative phosphorylation while achieving redox balance, of which antioxidant intervention achieves a low-level redox balance and moderate exercise achieves a high-level redox balance.

Our study, for the first time, found hepatic AMPK activation could act as a biomarker of dynamic redox balance during exercise to benefit glycaemic control in diabetic rats. Different intensities of exercise-induced ROS production could profoundly alter the cellular redox microenvironment and directly regulate the activity and expression of hepatic AMPK through a redox-related mechanism. Moderate exercise (including CE and IE) appropriately promotes oxidation in the liver, thus, compensatory promotes the level of reduction, forming a high level of redox balance. Excessive exercise producing ROS exceeding the redox threshold caused oxidative damage with a significant increase in MDA and urea nitrogen levels. Specifically, optimal ROS directly promoted AMPK activation via glutathionylation and upregulated glycolysis and aerobic oxidation in L02 cells, which was consistent with the animal experiments. However, excessive ROS inhibits the activity and expression of AMPK in L02 cells, which might be related with the oxidative stress induced protein degradation (Fig. 7). Accordingly, we speculate that the antioxidant intervention based on moderate exercise might offset the effect of exercise, but the antioxidant intervention after excessive exercise could restore redox balance. In addition, the number of mitochondria and the function of aerobic oxidation in the IE group was significantly higher than those in the CE group. In the EE group, the autophagy and fission of liver mitochondria are also up-regulated. These results indicate that the phosphorylation and expression of AMPK can act as a sensitive biomarker during exercise in diabetic rats, reflecting the threshold of redox balance and the range of exercise appropriateness.

**Fig. 7.**
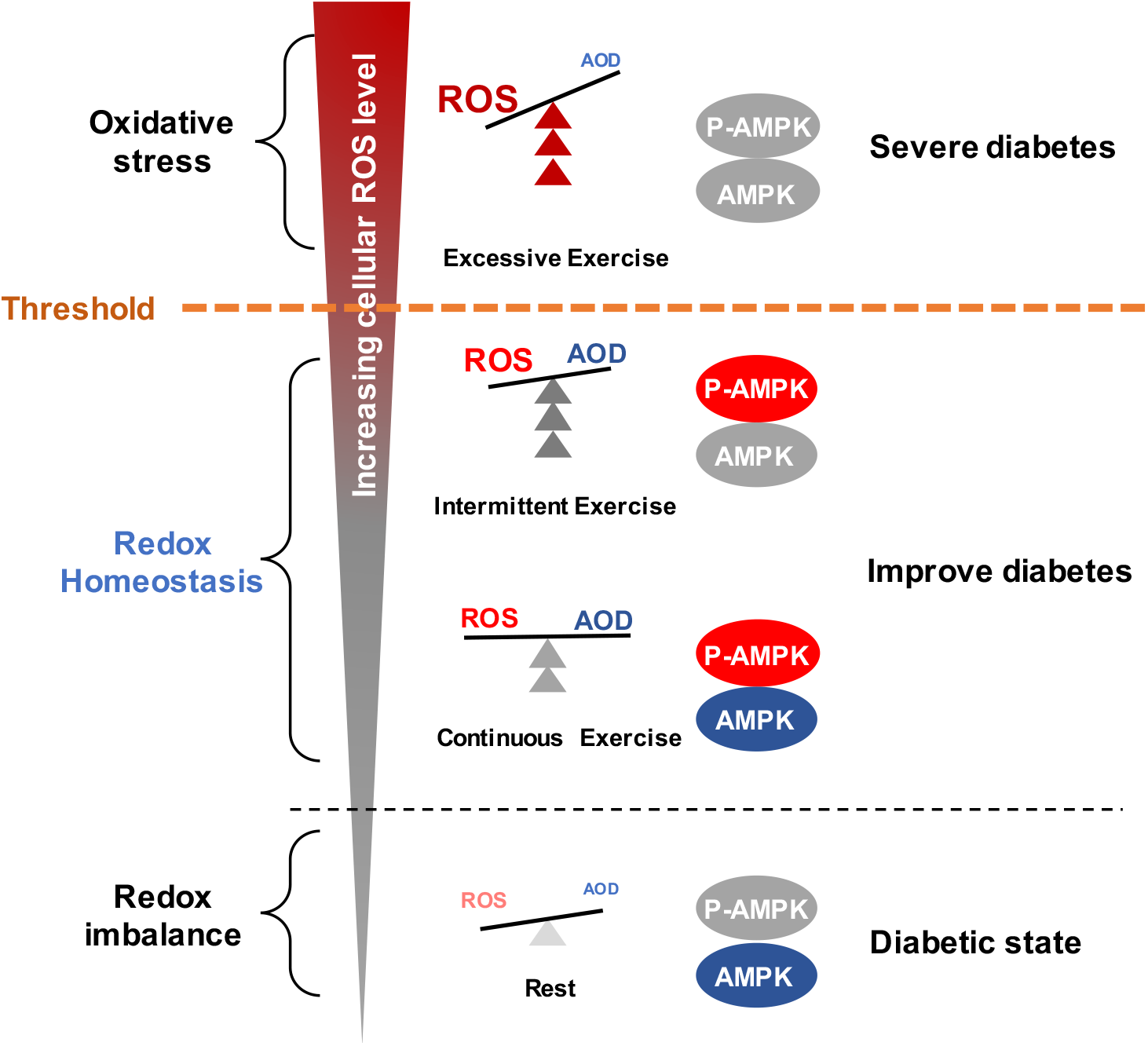
Schematic diagram of redox balance threshold. Moderate exercise promotes the activity of antioxidant enzymes by generating benign ROS, directly promotes AMPK-mediated glycolysis and aerobic oxidation, thus reduces glucose levels in blood and liver. Excessive exercise causes excess ROS and exceed the redox balance threshold, inhibiting AMPK activity and expression, thus leading to exacerbation of diabetes. (AMPK and P-AMPK in grey circles indicate decrease, red circles indicate increase, blue circles indicate no significant changes).

Together, our results illustrate the different regulatory mechanisms of exercise and antioxidant intervention on redox balance and blood glucose level in diabetes. Moderate exercise promoted hepatic ROS production and up-regulated a compensatory increase antioxidant capability, forming a high-level balance of redox state. In contrast, antioxidant intervention scavenged the hepatic free radical to form a delicate low-level balance of redox state. Moreover, excessive exercise led to redox imbalance due to excess ROS levels. Hepatic AMPK activation could act as a sign and hallmark of moderate exercise and dynamic redox balance to guide appropriate exercise or antioxidant intervention. These results illustrate that it is necessary to develop a moderate exercise program according to the REDOX microenvironment of diabetes patients and provide theoretical evidence for the precise regulation of diabetes by antioxidants and exercise.

## Materials and Methods

### 1. Materials

T-AOC kit was supplied by Changzhou Redox Biological Technology Corporation (Jiangsu, CN). Antibodies against Actin, Acetylated-Lysine, P-PFK2, Ace-SOD2, ATG5, LC3A/B, GAPDH, MFN1 and IgG-HRP were purchased from Cell Signaling Technology (USA). Antibodies against CAT, PRX1, AMPKa1, GRX1, GRX2, SOD2, HSP90, COX1, COX2 and PFK2 were purchased from ProteinTech (Wuhan, CN). Antibodies against 3-NT, 4HNE, NOX4 and PGC1-α were purchased from Abcam. Antibodies against p-AMPKα1/α2 were purchased from SAB (Signalway Antibody, USA). The detailed antibody information is shown in Supplementary Table S1. High-fat diet (HFD) were purchased from Shanghai SLRC laboratory animal Company Ltd (Shanghai, China) and the nutritional composition is shown in Supplementary Table S2.

### 2. Animal

Male SD rats (150–160 g body weight, 6-8 weeks) were purchased from Fudan University Animal Center (Shanghai, China). All animal care and experimental procedures were approved by the Fudan University Institutional Laboratory Animal Ethics Committee (NO. 20170223-123). Animals were housed in a pathogen free environment with 12 h dark/light cycles.

### 3. Establishment of diabetic rat model

Rats were divided into six groups in a non-blinded, randomized manner: Control (Ctl), STZ+HFD diabetic rat (T2D), Continuous exercise+STZ+HFD diabetic rat (T2D+CE), intermittent exercise+STZ+HFD diabetic rat (T2D+IE), excessive exercise+STZ+HFD diabetic rat (T2D+EE) and Apocynin+STZ+HFD diabetic rat (T2D+APO) (n=8 per group). The sample size was calculated according to the Power Curve. The diabetic rats model was established by 12hr-fasting followed by intraperitoneal injection of 0.1 M streptozotocin (STZ) citrate solution (pH 4.5) at a dose of 35 mg/kg for Day1, and 35 mg/kg for Day 2 at the 5^th^ week. The HFD was started from the 1^st^ week to the 8^th^ week. After 8 weeks of intervention, the mice were sacrificed. The tissues and plasma were collected and preserved at ™80 °C for further analysis.

Rat were acclimated to treadmill running for 3 days before the initiation of the experiments and the exercise training intervention was continued for 4 weeks (5 times per week). All animals were randomized before the initiation of exercise tests.

#### Continuous exercise

The initial speed was 15 m/min, and the speed was increased by 3 m/min every 5 min. After the speed reached 20 m/min, the speed was maintained for another 60 min with slope of 5%. The exercise intensity was 64%-76% VO_2max_ (*28, 61, 62*).

#### Intermittent exercise

The initial speed was 15 m/min, and the speed was increased by 3 m/min every 5 min. After the speed reached 20 m/min, the speed was maintained for 20 min and then 5 min rest at 5 m/min. The training was continued for 3 times, and the total running time is 60 min with two 5 min rest with slope of 5%.

#### Excessive exercise

The initial speed was 15 m/min, and the speed was increased by 3 m/min every 5 min. After the speed reached 50 m/min, the speed was maintained for another 60 min with slope of 5%. The exercise intensity was higher than 80% VO_2max_.

OGTT was performed in the fasting mice with intraperitoneal injection of glucose at 1 g/kg of body weight, and glucose was measured at 15min, 30 min, 60 min and 120 min, respectively. Blood glucose was determined by glucometer (Roche, Switzerland).

### 4. Cell culture

Normal Human Hepatic Cell Line L02 cells (Cell Bank of Chinese Academy of Sciences) were grown in DMEM supplemented with 10% FBS (GIBCO, USA) in a humidified incubator (Forma Scientific) at 37 °C and 5% CO_2_ as described previously. The medium were supplemented with 10% FBS (GIBCO, USA), 2 mmol/l glutamine, 1 mmol/l sodium pyruvate, 10 mmol/l HEPES, 50 μmol/l β-mercaptoethanol, 105 U/l penicillin and streptomycin. Glutamine and sodium pyruvate were purchased from Sinopharm Chemical Reagent Co., Ltd, HEPES were purchased from Beyotime Biotechnology (Shanghai, CN). All cell lines used in the study were tested for mycoplasma and were STR profiled.

### 5. Flow cytometry

For measurement of intracellular Superoxide and H_2_O_2_, hepatocytes were stained with 5 μM hydroethidine (superoxide indicator) and 10 μM H_2_ DCFDA (Thermo Fisher Scientific, USA). Stained cells were analyzed with NovoCyte Quanteon flow cytometer (Agilent Technologies, Inc.), and acquired data were analyzed with NovoExpress software (Agilent Technologies, Inc.) and FlowJo software (TreeStar, Ashland, OR).

### 6. ATP and AMP content analysis

Liver tissue (20–30 mg) were homogenized on ice by perchloric acid. Homogenized samples were centrifuged for 12,000 rpm at 4 °C (30 min). Supernatant was then neutralized with 4 M K2CO3, followed by further centrifugation for 12,000 rpm at 4 °C for 20 min. Supernatant was obtained for the determination of ATP and AMP content by high performance liquid chromatography (HPLC). The detection wavelength was 254 nm.

### 7. Metabolite profiling detection

Cellular metabolites were extracted and analysed by LC-MS/MS. Ferulic acid was added as an internal standard to metabolite extracts, and metabolite abundance was expressed relative to the internal standard and normalized to cell number. Mass isotopomer distribution was determined by LC-MS/MS (AB SCIEX Triple-TOF 4600) with selective reaction monitoring (SRM) in positive/negative mode

### 8. Western blot analysis

Cells and tissues were lysed in a buffer containing 1% Nonidet P-40, 0.25% sodium deoxycholate, 150 mmol/l NaCl, 10 mmol/l Tris, 1 mmol/l EGTA, 1% proteinase and phosphatase inhibitor cocktails (Sigma-Aldrich) at 4 °C for 30 min. Cell lysates were resolved by sodium dodecyl sulfate-polyacrylamide (SDS-PAGE) gel electrophoresis, transferred to polyvinylidene fluoride (PVDF) membranes, and immunoblotted with primary antibodies. Membranes were incubated with HRP-conjugated secondary antibodies and visualized using chemiluminescent substrate (ECL; Tanon, CN) and Tanon-5200 Chemiluminesent Imaging System (Tanon, CN).

### 9. Transmission electron microscope (TEM)

Rat liver tissue (1 mm*1 mm) were fixed by paraformaldehyde. The samples were examined with a Jeol Jem-100SV electron microscope (Japan) which was operated at 80 Kv after fixed by 3% glutaraldehyde in 0.1 M phosphate buffer (pH 7.3) at Institute of Electron microscopy, Shanghai Medical College of Fudan University.

### 10. Statistics

The experimental data were expressed as mean ± SEM. One-way ANOVA was used to compare among groups. Data analysis was conducted by Graphpad prism 9 statistical analysis software. p < 0.05 was considered statistically significant. Data are expressed as means ± SEM; n = 3 for cells experiment (n = 3 represents three times of individual experiment); n = 8 for animal experiment.

## Acknowledgments

The authors thank Dr. Xiaodong Zhang from Chengdu Brilliant Pharmaceuticals for his proof reading and editing of the manuscript. The authors are also indebted to Dr. Rutan Zhang and Prof. Liang Qiao from Fudan University, for analysis of LC-MS/MS data. The authors thank Mr. Yipei He, Dr. Xiao Zhang, Mrs Lihan Jiang, Mr. Kelei Dong for their assistance in animal experiments. The authors are also indebted to Institute of Electronmicroscopy from Fudan University, for the help on electronmicroscopy analysis.

This work was supported by grants from the National Natural Science Foundation of China (Grants No. 31770916).

## Author contributions

M.W., A.Z., C.Y., X.L., C.Z. and Q.L. conducted experiments. L.S. and D.S designed the experiments and analyzed data. Y.F. and F.X. provided professional consultation on exercise mode. H.G. provided professional suggestion on electronmicroscopy analysis. L.S. and D.S. supervised the project and wrote the manuscript.

## Data and materials availability

All data are available in the main text or the supplementary materials.

## Graphic Abstract

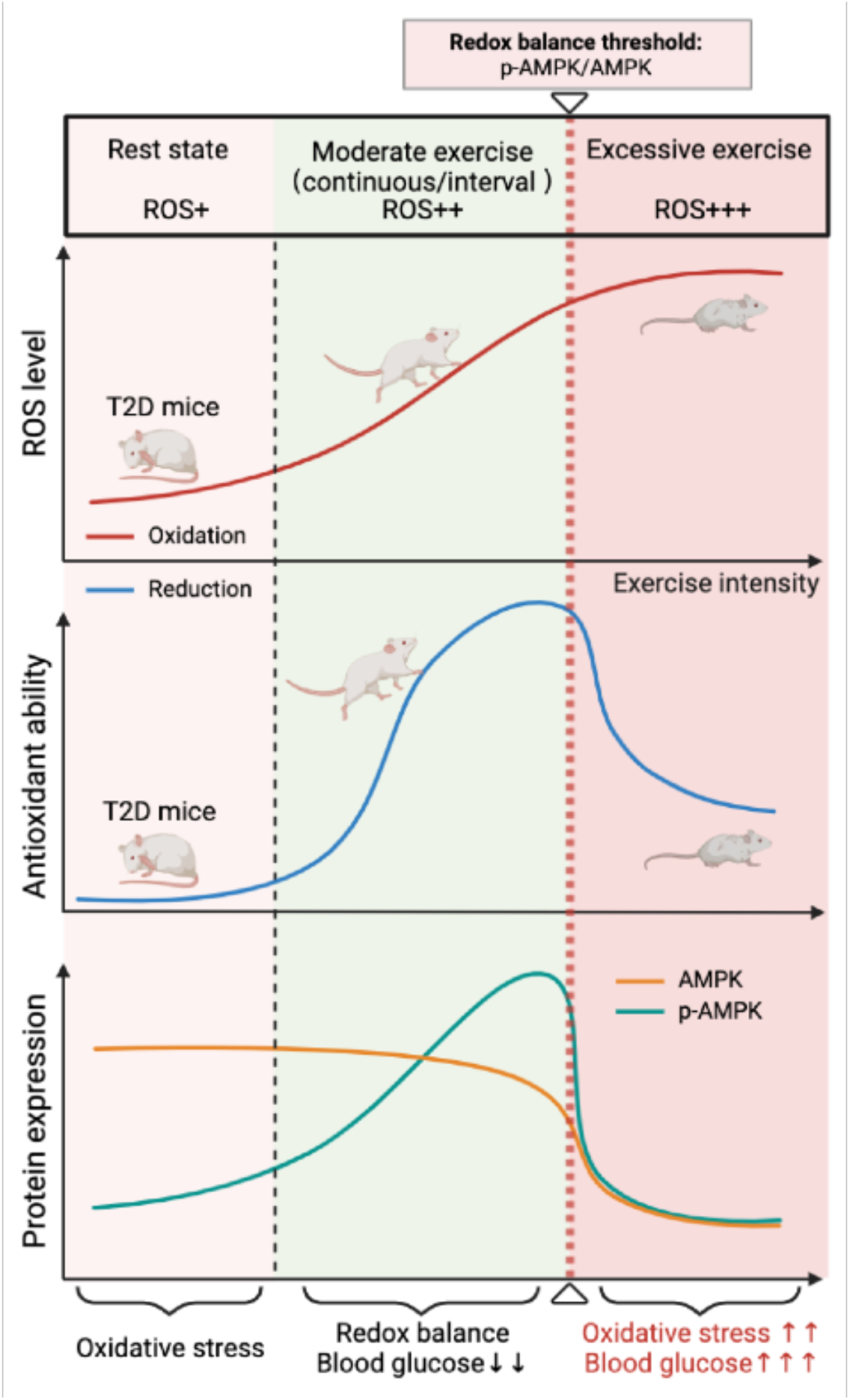

