## Supplemental Table for "Hepatic AMPK activation in response to dynamic REDOX balance is a biomarker of exercise to improve blood glucose control"

^4^ Clinical Nutrition Center, Huadong Hospital Affiliated to Fudan University, Shanghai, 200032, People's Republic of China

**Supplementary table:** Antibody information

|  | Name | Supplier | Cat No. |
| --- | --- | --- | --- |
| 1 | CAT | ProteinTech (Wuhan, CN) | 21260-1-AP |
| 2 | PRX1 | ProteinTech (Wuhan, CN) | 15816-1-AP |
| 3 | AMPKa1 | ProteinTech (Wuhan, CN) | 10929-2-AP |
| 4 | GRX1 | ProteinTech (Wuhan, CN) | 15804-1-AP |
| 5 | GRX2 | ProteinTech (Wuhan, CN) | 13381-1-AP |
| 6 | SOD2 | ProteinTech (Wuhan, CN) | 24127-1-AP |
| 7 | HSP90 | ProteinTech (Wuhan, CN) | 13171-1-AP |
| 8 | COX1 | ProteinTech (Wuhan, CN) | 13393-1-AP |
| 9 | COX2 | ProteinTech (Wuhan, CN) | 12375-1-AP |
| 10 | PFK2 | ProteinTech (Wuhan, CN) | 17838-1-AP |
| 11 | P-AMPKa1/a2 | SAB | #11183 |
| 12 | 3-NT | Abcam | ab61392 |
| 13 | 4HNE | Abcam | ab46545 |
| 14 | NOX4 | Abcam | Ab133303 |
| 15 | PGC1-α | Abcam | ab54481 |
| 16 | Actin | Cell Signaling Technology (USA) | #3700 |
| 17 | Acetylated-Lysine | Cell Signaling Technology (USA) | #9441 |
| 18 | P-PFK2 | Cell Signaling Technology (USA) | #13064 |
| 19 | Ace-SOD2 | ProteinTech (Wuhan, CN) |  |
| 20 | ATG5 | Cell Signaling Technology (USA) | #12994 |
| 21 | LC3A/B | Cell Signaling Technology (USA) | #12741 |
| 22 | Anti-rabbit IgG-HRP | Cell Signaling Technology (USA) | #7074 |
| 23 | Anti-mouse IgG-HRP | Cell Signaling Technology (USA) | #7076 |
| 24 | GAPDH | Cell Signaling Technology (USA) | #2118 |
| 25 | MFN1 | Cell Signaling Technology (USA) | #14739 |
| 26 | Fis1 | Merck | ABC67 |
